# MeCP2 deficiency exacerbates the neuroinflammatory setting and autoreactive response during an autoimmune challenge: implications for Rett Syndrome

**DOI:** 10.1101/2020.08.05.238683

**Authors:** MI Zalosnik, MC Fabio, ML Bertoldi, CL Castañares, AL Degano

## Abstract

**Background:** Rett syndrome is a severe and progressive neurological disorder linked to mutations in the MeCP2 gene located on the X chromosome. So far it has not been established how the presence of a mutant form of MeCP2 can maintain essential regulation of immune responses to support the normal homeostasis of individuals. Since MeCP2 is mostly expressed as a “partially functional” protein in humans with RTT, the aim of our work was to evaluate whether a mutation in MeCP2 interferes with the induction of neuroinflammatory responses in real time.

**Methods:** We used MeCP2^308/y^ mouse model (MUT) and exposed it to an autoimmune challenge, experimental autoimmune encephalomyelitis (EAE). WT and MUT mice were immunized with CFA-MOG or CFA alone (control) and clinical scores were evaluated daily. Animals were sacrificed at either 12 days post-induction (dpi, acute stage) or 30 dpi (chronic stage) and spleen and spinal cord were collected from individual mice for further studies. Cellular infiltration and microgliosis was evaluated by IHC. Cytokine production was assessed in spinal cord and in cultured splenocytes after MOG activation ex-vivo by cytometry and real time RT-PCR.

**Results:** Our results showed that MeCP2 deficiency increased the susceptibility to develop EAE, along with a defective induction of anti-inflammatory responses and an exacerbated MOG-specific reactivity with high IFNγ expression in peripheral immune sites. During the chronic stage, an increase in gene expression of pro-inflammatory cytokines (IFNγ, TNFα and IL-1β) and downregulation of genes relevant for immune regulation (IL-10, FoxP3 and CX3CR1) was found in MUT-EAE spinal cords.

**Conclusions:** This is the first study performed in a MeCP2 mutant mouse model that explores the pathophysiology and neuroinflammation in the context of an autoimmune challenge. We could establish that an MeCP2 mutation act intrinsically affecting neuroimmune interactions by promoting an inflammatory environment and a deficient immune regulatory setting. These results are relevant for understanding the consequences of MeCP2 mutations on immune homeostasis in MeCP2-related disorders, as well as setting the bases for further therapeutic interventions that consider the immune status in patients.

## INTRODUCTION

Methyl-CpG binding protein 2 (MeCP2) is an X-chromosome nuclear protein that recognizes and binds specifically to methylated cytosine residues in the DNA, in regions enriched with adjacent A/T bases [1]. MeCP2 has been reported to play multifunctional roles in regulating gene expression, participating in RNA splicing [2], chromatin remodeling [3], [4], transcriptional activation [5] and repression [6]. This protein is expressed in all tissues although it is particularly abundant in the central nervous system (CNS) [7]. Since loss-of-function mutations in the *mecp2* locus is the etiological cause of 95% of typical Rett syndrome [5], [8], MeCP2 role in the CNS has been extensively studied. Rett syndrome (RTT, MIM 312750) is one of the most common causes of mental retardation in females, with an incidence of 1 in 10,000–15,000 [8]. Classical RTT patients appear to develop overtly normal until 6–18 months of age, followed by the onset of symptoms, which include gradual loss of speech and purposeful hand use, stereotypic hand movements, seizures, autism, ataxia, and breathing disturbances [9].

Although most studies have explored the critical role of MeCP2 in neural function, MeCP2 expression is ubiquitous and the pathogenesis of RTT compromise other systems besides the CNS [10]. In this regard, polymorphisms in *MECP2* have been linked to increased susceptibility to autoimmune diseases in humans, such as systemic lupus erythematosus (SLE) [11], [12], thyroid diseases [13] and primary Sjogren’s syndrome [14]. MeCP2 is expressed in immune cells, and alterations in MeCP2 expression levels affect immune function and cytokine production [15], [16]. In particular, it has been proposed that dysfunctions in myeloid cells such as macrophages and microglia, would actively contribute to the pathogenesis of RTT. In RTT patients, abnormal levels of macrophage-related cytokines, such as TNFα, IL-6, IL-12p70, IL-10 and TGF-β1 have been identified [17]–[19]. Using *Mecp2*-null mice, it was determined that microglial populations decreased in number as the disease progressed, and these cells displayed defective immune activation [20]. Moreover, these authors showed that wild-type microglia introduced by bone marrow transplantation, ameliorated some of the deficits observed in *Mecp2*-null mice. However, MeCP2 expression is also important in other immune cells. MeCP2 seems crucial to regulate naive CD4^+^ T cells differentiation to T helper type 1 (Th1) and Th17 cells[21], and it is also required to maintain immune tolerance by supporting the stable expression of FoxP3 expression, a protein involved in the development and function of regulatory T cells [22].

All these data suggest that M*ecp2* mutations may affect the immune homeostasis and thus, it is reasonable to consider that the immune system may play an active role in the generation and/or maintenance of RTT alterations. However, there is no clear consensus about which pathways or genes are regulated *in vivo* by MeCP2 in the context of immune activation. Another limitation of previous studies, is that they used “null” mouse models, with total absence of MeCP2 protein. From a potentially translational point of view, the interpretation of these results is complex in the context of RTT and associated disorders, in which MeCP2 protein is indeed expressed, although at lower levels or as a functionally deficient protein.

In the present work we used a well characterized RTT mouse model (MeCP2^308/y^), that carries an early termination codon and generates an MeCP2 protein truncated at amino acid 308, with loss of the C-terminal region [23]. This truncated form of MeCP2 lacks several regulatory sites, such as phosphorylation sites that are key for co-repressors/co-activators binding [24], [25]. Likewise, the absence of the C-terminal domain leads to lower stability and shorter half-life of the truncated protein compared to the wild-type (WT) form [26]. As a consequence, this model represents a condition of “MeCP2 function deficiency”, as opposed to “total MeCP2 absence” represented by *Mecp2*-null models. The MeCP2^308/y^ mutant model shows a progressive neurological phenotype and longer lifespan, making it more suitable for immune studies.

Preliminary studies in our lab assessed basal levels of autoantibodies against CNS and myelin proteins in sera from both MeCP2^308/y^ and *Mecp2*-null mice. We found no differential auto-reactivity patterns in these mouse models respective to their WT littermates (data not shown), suggesting that immune activation would be important for MeCP2 regulation of immune responses. Thus, in the present work we set to characterize the role of MeCP2 during the progression of Experimental Autoimmune Encephalomyelitis (EAE), considering that this experimental model gives us the opportunity to characterize the autoimmune response and concomitant neuroinflammatory status. Our results showed that MeCP2 deficiency increased the susceptibility to develop EAE, along with an exacerbated inflammatory profile and a defective induction of anti-inflammatory responses (both in peripheral immune sites and in the CNS). Importantly, in the absence of immune activation, we detected no differences between WT and MeCP2^308/y^ in any of the parameters tested. To our knowledge, this is the first report that explored the role of MeCP2 in the pathophysiology and neuroinflammatory response in the context of an autoimmune challenge. Our results suggest that MeCP2 acts intrinsically upon immune activation, affecting neuroimmune homeostasis by regulating the pro-inflammatory/anti-inflammatory balance *in vivo*.

## METHODS

### Animals

For our studies we used the Mecp2^308/y^ mouse model (B6.129S-Mecp2 /J, Stock 005439, The Jackson Labs). These animals carry a premature stop codon at amino acid 308, generating a truncated MeCP2 protein that lacks the C-terminal region [23]. The animals were kept at the facility of the “Departamento de Química Biológica-CIQUIBIC, Facultad de Ciencias Químicas” (Universidad Nacional de Córdoba, Argentina) and maintained in a C57BL/6J background. All the experiments were performed using only hemizygous MeCP2 males (MeCP2 MUT) and their corresponding WT male littermates as control, in order to avoid the high variability caused by random X-inactivation in females. Animal procedures were fully reviewed and approved by the local animal committee (School of Chemistry, National University of Córdoba, Protocol 2015-832, renewed as 2019-2879), which follows guidelines from the National Institute of Health. Every effort was made to minimize both the number of animals used and their suffering. All the animals were housed and maintained on a 12/12 h light/dark cycle (lights on at 7 am) with food and water *ad-libitum*. 2 weeks old mice were genotyped following the protocol provided by The Jackson Labs (https://www.jax.org/protocol/search) and then male littermates were divided into 4 different experimental groups: MeCP2-wildtype animals treated with complete Freund’s adjuvant (<u>WT-CFA</u>) or immunized with MOG (<u>WT-EAE</u>); MeCP2-mutant animals treated with complete Freund’s adjuvant (<u>MUT-CFA</u>) or immunized with MOG (<u>MUT-EAE</u>).

### EAE induction

Male MeCP2 WT and MUT mice, 9 weeks old, were anesthetized via i.p. with a mixture of xylazine and ketamine (16 mg/kg and 80 mg/kg respectively) and immunized subcutaneously at the right and left flanks with 200μl of an emulsion containing 200μg of myelin oligodendrocyte glycoprotein peptide (MOG_35-55_, NH_2_-MEVGWYRSPFSRVVHLYRNGK-COOH; synthesized at the Johns Hopkins University Synthesis & Sequencing Core Facility, Baltimore, MD, USA). The peptide was dissolved in sterile water at 2 mg/ml, mixed at a 1:1 ratio with complete Freund’s adjuvant (CFA, Sigma-Aldrich Co., St. Louis, MO, USA), supplemented with 4 mg/ml of *Mycobacterium tuberculosis*. Pertussis toxin (200 ng List Labs, USA) was dissolved in 100 μl of phosphate-buffered saline (PBS) and injected i.p. the same day of the immunization and 48 h later. Mice were scored daily for EAE symptoms starting at 6 dpi. They were euthanized at either, 12 dpi (acute stage) or at 30 dpi (chronic stage). Given that the clinical signs of the tail and each leg did not develop together, we graded the signs for tail and hind legs separately and establishing a scale of 0-8, where 2 corresponds to the maximum tail score and 3 the maximum for each leg. The tail abnormalities were scored as follows: 0, no deficits, (tail moves and can be raised, tail wraps around a round object if mouse is held at the base of the tail); 1 partial loss of tail tone, 2 total tail paralysis. Each hind limb was graded according to the following scale: 0, normal gait; 1, mild hind limb weakness; 2, dragged limp with abnormal gait; 3, complete hind limb paralysis with no residual movement. Although the MeCP2 308 mutant mice could show altered motor skills after 8 weeks of age, mutant mice injected with CFA showed no differences in weight compared with WT, no obvious symptoms or motor abnormalities during the period tested (up to 30 dpi, i.e. 13 weeks old mice).

### Tissue processing

Animals were deeply anesthetized with an i.p. injection of ketamine-xylazine cocktail (16 mg/kg and 80 mg/kg respectively) and perfused intracardially with ice-cold PBS and then with ice-cold 4% paraformaldehyde in PBS. Spinal cords were removed and placed in 4% paraformaldehyde for 2 more hours. Then the tissues were buffered first, in 15% sucrose overnight and then in 30% sucrose solution for another 24 h at 4°C. Finally, the tissues were embedded in mounting medium Cryoplast (Biopack, Buenos Aires, Argentina) and stored at −80°C until sectioning. 20 μm-thick serial sections of lumbar spinal cord were obtained using a cryostat (CM1510 S model, Leica Microsystems, Wetzlar, Germany), attached to aminopropyltriethoxysilane (APES, Sigma #A3648) treated-glass slides, and stored at −80°C until use.

### Toluidine Blue Staining

Six-step sections (20 μm) of lumbar spinal cords were incubated in toluidine blue 0.25 % (ANEDRA, Tigre, Buenos Aires, Argentina) diluted in acid buffer (0.088 M acetic acid, 0.012 M sodium acetate) during 30sec and then washed in distilled water. Slides were left to dry and mounted using DPX (Sigma-Aldrich Co., St. Louis, MO, USA). Digital images were collected under optical microscope using a 10X and 20X objective. Slides were assessed in a blinded fashion for infiltrating immune cells in spinal cord in different anatomical compartments (meninges and parenchyma). The level of infiltrating cells was scored for the meninges: 0, no infiltrates; 1, few isolated cells; 2 cell focuses; 3 extensive cell focuses. For parenchyma: 0, no infiltrates; 1, one or two cell focus; 2, more than 2 small focuses or one extensive focus; 3, more than two deep or extensive focus of cells. For each animal, 6 serial coronal sections of lumbar spinal cords were analyzed and the infiltration level was obtained as the sum of the score infiltration in each of the sections.

### Immunofluorescence

Six-step serial coronal sections from lumbar spinal cords (20 μm) were blocked in Blocking Buffer (4% bovine fetal serum, 0,3% Triton in PBS) for 60 min at room temperature. Primary antibody (anti-Iba-1, 1:500, Abcam) was diluted in blocking buffer and incubated overnight at 4°C in a humid light-tight box. Next, after 2 washes in PBS, the slides were incubated with the secondary antibody (Alexa 488-conjugated anti-rat IgG, 1:500) diluted in PBS for 1 h at room temperature. Afterwards, slides were incubated in DAPI for 5 min, washed 3 times in PBS and mounted using Mowiol 4-88 reagent (Aldrich, St. Louis, MO, USA). Images were acquired using a FV1000 confocal microscope (Olympus, Tokyo, Japan) equipped with argon/helium/neon lasers and 20X and 40X objectives. For each animal, 6 serial coronal sections of lumbar spinal cords were obtained and 5 fields per each coronal section was photographed in Z-projection using a 40X objective. The obtained images were subjected to a threshold and processed, to analyze the positive area of Iba expression (Iba-1^+^ area), using a size-based particle exclusion plugin (software ImageJ).

### Real Time RT-PCR

Gene expression analysis was performed by real time RT-PCR. Mice were euthanized by cervical dislocation and lumbar spinal cords were removed. Tissues were homogenized and resuspended in 1 ml of TRIzol reagent (Invitrogen, Carlsbad, CA, USA) and then we proceeded according to manufacturer’s instructions for RNA extraction. 2 μg of RNA was incubated at room temperature for 15 min with DNase I (Invitrogen, Carlsbad, CA, USA). The product was incubated with random hexamer primers, deoxynucleotides and the reverse transcriptase M-MLV (Moloney Murine Leukemia Virus Reverse Transcriptase), all from Promega, Madison, WI, USA. Reverse transcription was performed following the manufacturer’s specifications, employing a thermocycler Mastercycler gradient (Eppendorf, Hamburg, Germany) in one cycle as follows: 6 min at 25 °C, 60 min at 37 °C, 18 min at 70 °C and 10 min at 4 °C. The generated cDNA was diluted with sterile milliQ water. For real time PCR reactions, 6 μl of cDNA was mixed with 0.375 μl of a 10 μM solution of each primer (sequences available upon request), 7.5 μl of 2× SYBR Green PCR Master Mix (Promega, Madison, WI, USA) and sterile milliQ water to a final volume of 15 μl per tube. Duplicates were prepared for each sample. Real-time PCR was performed on the thermal cycler Rotor-Gene Q (Qiagen, Venlo, Limburg, Netherlands) according to the following protocol: Initial denaturation 10 min at 95 °C, amplification (45 cycles) with denaturation 15 s at 95°C, annealing 30 s at 60°C and extension 30 s at 70°C. To confirm the presence of a single product, agarose electrophoresis and a melting curve of the DNA was always made covering the range of 50-95°C. Relative gene expression levels were quantified using the comparative ΔΔCT method [27], [28]. This method normalized CT values of the detected gene to the average of GAPDH and calculated the relative expression values as fold changes of the control group (WT-CFA), which was set at 1.

### Splenocytes culture and cytokine analysis

Control and EAE mice were euthanized 12 days post EAE induction (dpi; acute phase) or at 30 dpi (chronic phase). Under sterile conditions, the spleen was dissected and kept at 4°C in sterile D-PBS with 2% endotoxin-free fetal bovine serum (FBS) and Gentamicin (50 μg/ml) until they were processed. After removing the fat, the spleen was disintegrated using a metal mesh. Subsequently, cells were incubated with 6 mL of red blood cell lysis buffer [Ammonium Chloride (NH4Cl) 0.15M, Potassium Carbonate (KCO_3_) 10mM, EDTANa_2_ 0.1 mM at pH 7.4] for 6 minutes and centrifugated for 10 min at 1,600 rpm at 4°C. The supernatant was discarded and the pellet was resuspended with D-PBS supplemented with 10% FBS and Gentamicin, and centrifuged for 10 min at 1600 rpm at 4°C. This last step was repeated twice. The precipitate containing the splenocytes was resuspended in complete RPMI 1640 medium [10% FBS, Gentamicin (50μg/mL), L-Glutamine (2mM). The number and viability of cells were estimated by counting in 0.2% trypan blue solution in Neubauer chamber. Splenocytes concentration was adjusted to 1 × 10^6^ cells/ml, cultured in complete RPMI medium and stimulated with MOG (1 μg/ml) or with complete medium alone, as negative control. Supernatants were collected at 72 h and levels of IFN-γ, TNF-α, IL-2, IL-4, IL-6, IL-10 and IL-17 were analyzed using the Cytometric Bead Array (CBA) Mouse Th1/Th2 Cytokine Kit (BD Biosciences) according to manufacturer’s instructions.

### Data analysis

The results are expressed as the mean ± SEM. Whenever possible, independent variables were analyzed using test t-student or two-way analysis of variance (ANOVA). Whenever ANOVA indicated significant effects (p≤0.05), a Tukey *Post-Hoc* test was carried out. In all cases, the assumptions of the analysis of variance (homogeneity of variance and normal distribution) were attained. In all statistical analysis, a p<0.05 was considered to represent a significant difference between groups. All the analyses were performed using the software GraphPad Prism version 6.0 (La Jolla, California USA).

## RESULTS

### Mecp2^308/y^ mice show early onset and more severe EAE clinical manifestations

In order to evaluate the role of MeCP2 during an immune challenge *in vivo*, EAE was induced in WT and MUT mice and clinical signs were evaluated daily. None of the mice died as a result of EAE during acute or chronic stage. In addition, no animals treated with CFA (WT or MUT) developed the disease or showed changes in their motor skills after EAE induction.

Daily clinical signs analysis showed significant differences in clinical scores between these two groups (Two-factor ANOVA with repeated measures. Genotype x EAE, p=0.0035). Between 8 dpi and 14 dpi, MUT-EAE showed more severe clinical signs than WT-EAE (Fig. 1.a). Importantly, MUT-EAE mice showed an early onset of the disease compared to WT-EAE (Fig. 1.b). In addition, the maximum clinical score (MSC) and disease index (DI) were calculated in order to further examine the severity in each experimental group. Although no significant differences were found in the MCS between WT and MUT animals (Fig. 1.c), the accumulated disease index was significantly higher in MUT mice (Fig. 1.d).

**Fig. 1.**
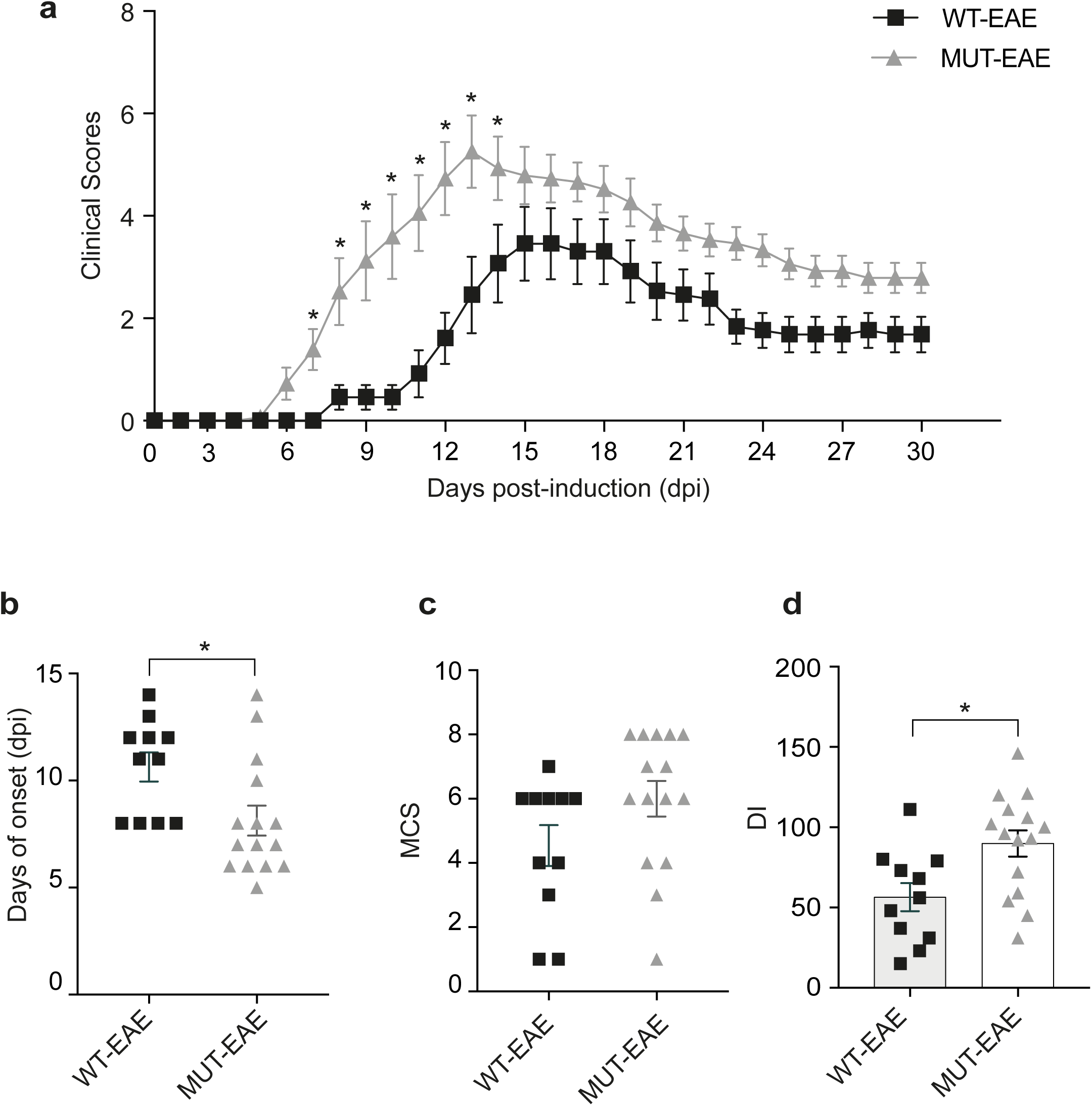
Clinical evaluation of EAE. **a**. WT and MeCP2 mutant mice (MUT) were immunized with MOG and clinical signs were evaluated daily for 30 days. MUT-EAE mice showed the first clinical signs between 6 and 8 dpi and more severe clinical signs throughout the acute and chronic stages with respect to WT. A two-factor ANOVA with repeated measures was performed. *p<0.05. n=13-15 per experimental group. **b-d**. Graphs summarizing the indexes calculated from the clinical information obtained. **b** DO (day of onset) post induction day in which the first clinical signs were observed. **c**. MCS (maximal clinical score) is the sum of the highest clinical score achieved by each mouse during the acute stage of the disease, divided by the number of mice that became ill in that group. **d**. (DI) Cumulative disease index is the sum of all clinical scores of each animal divided by the total number of animals in that experimental group. It was determined that MUT-EAE mice showed higher incidence of disease, higher rate of accumulated disease and early onset of EAE. All values are presented as media ± SEM. For all indices, statistical analysis was performed using the t-Student test. *p<0.05. n=13-15 per experimental group.

It should be noted that the incidence of the disease was 100% for MUT mice, while for WT it was only 84%; WT mice that did not develop EAE were discarded from subsequent analyzes. In addition, during the chronic stage, MUT-EAE mice had a slower recovery at the level of motor condition although by 30 dpi, both groups showed similar clinical scores. These results suggest that MeCP2-mutant animals have a greater predisposition or vulnerability to develop EAE, as they showed an early onset of the disease, greater severity of clinical signs and required more time for partial recovery in chronic phase.

### Splenocytes from Mecp2^308/y^ mice display a pro-inflammatory profile after MOG recall stimulation ex vivo

The initial immune response and maintenance of EAE occur mainly at the level of the lymphatic organs, where the antigen presentation triggers the activation and proliferation of effector cells [29]. Therefore, in order to understand the differences observed in EAE progress between WT and MUT animals, we set to examine the immune profile and the specific response to MOG. To that end, splenocytes were isolated from WT and MUT animals either in EAE acute or chronic stage and were re-stimulated with MOG *in vitro* (+MOG). After 72 hours of incubation, supernatants from each condition were isolated and concentrations of different cytokines (IFNγ, TNFα, IL6, IL17, IL2, IL4, IL10) were determined using a specific kit for Th1 / Th2 / Th17 cytokines (as shown in methods).

#### Acute stage

We found no detectable levels of the tested cytokines in supernatants of cultured splenocytes isolated from CFA animals, regardless their genotype and *in vitro* re stimulation with MOG (data not shown). While similar levels of MOG-induced pro-inflammatory cytokines (IFNγ, TNFα, IL17 and IL6) were secreted by WT-EAE and MUT-EAE splenocytes (Fig. 2a-d), significant differences were found for IL-2, IL-10 and IL-4, when considering the interaction between genotype and MOG stimulation (Fig. 2e-g). In this sense, MUT-EAE splenocytes +MOG, responded by increasing the production of IL-2 and IL-4, which was significantly different from the WT-EAE +MOG group. On the other hand, we observed a lower production of IL-10 from MUT-EAE +MOG splenocytes compared to WT-EAE +MOG group.

**Fig. 2.**
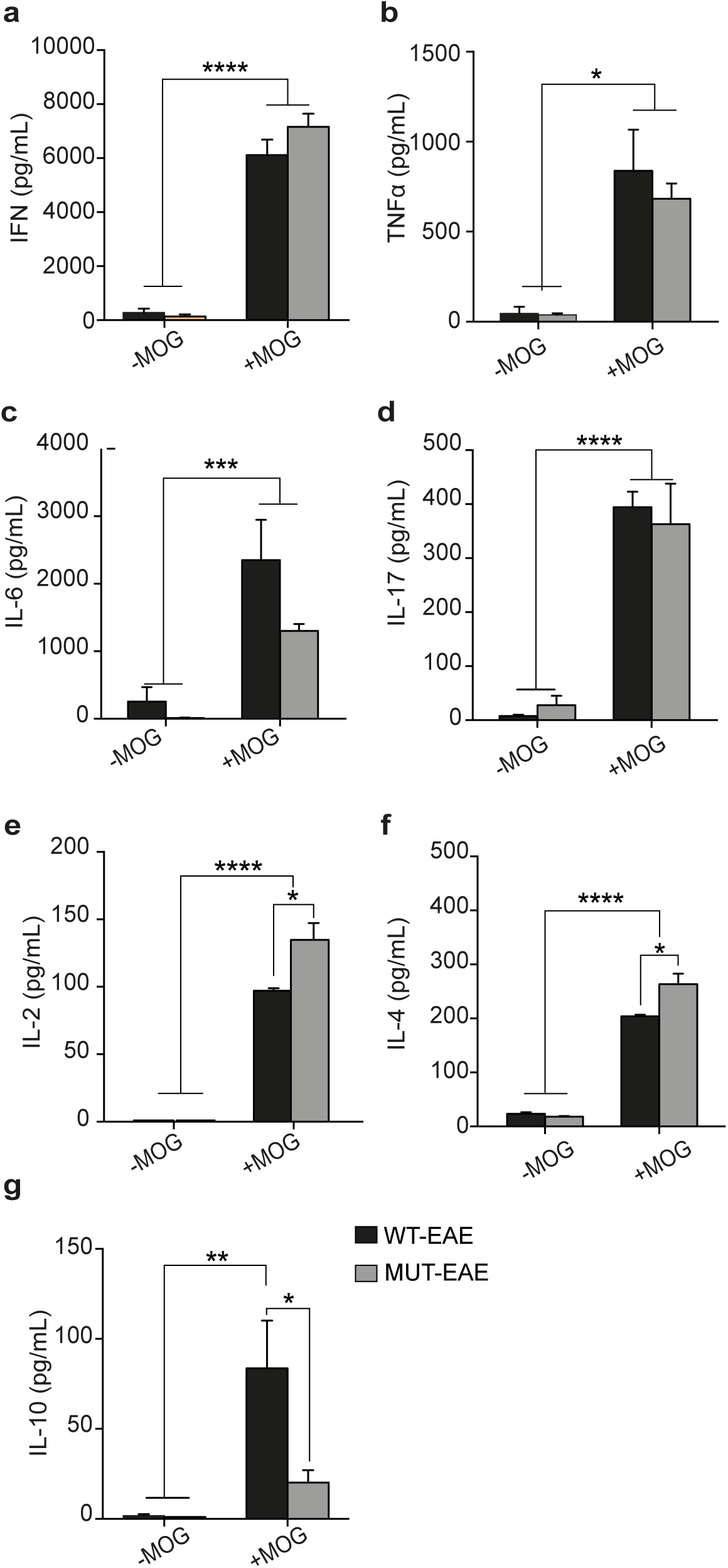
Cytokine production by splenocytes isolated during acute stage of EAE. The production of cytokines in supernatants by splenocytes isolated from WT-EAE and MUT-EAE animals was assessed in the absence or presence of MOG for 72 hours using a cytometry kit (-MOG, +MOG). For all the cytokines analyzed: **a**. IFNγ **b**. TNFα **c**. IL-6 **d**. IL-17, **e**. IL-4, **f**. IL-2 -except for **g**. IL-10, the *in vitro* re-stimulation with MOG showed a significant increase in production compared to the absence of MOG in culture. IL-2 and IL-4 production by MUT-EAE +MOG splenocytes was higher compared to WT-EAE +MOG, while IL-10 in the latter group was significantly higher. All values are presented as media ± SEM. Two-way ANOVA and Tukey post hoc test performed. *p<0.05; **p<0.0021 *** p<0.0002; ****p<0.0001. n = 3-5 per experimental group.

Considering that the progression of immune response along EAE clinical course, depends on the balance of pro- and anti-inflammatory cytokines, we calculated Th2/Th1 ratios between measured concentrations of anti-inflammatory cytokines (IL-4 and IL-10) and pro-inflammatory cytokines (IFNγ and TNFα) secreted by WT-EAE and MUT-EAE splenocytes in response to MOG (Fig. 3). No significant differences were found in IL-4/TNFα or IL-4/IFNγ ratios between WT and MUT groups (Fig. 3a). However, IL-10/IFNγ ratio was significantly lower and IL-10/TNFα ratio appeared also reduced compared to WT-EAE group (Fig. 3b). Thus, splenocytes from MUT-EAE mice responded to MOG generating a skewed Th1 proinflammatory profile.

**Fig. 3.**
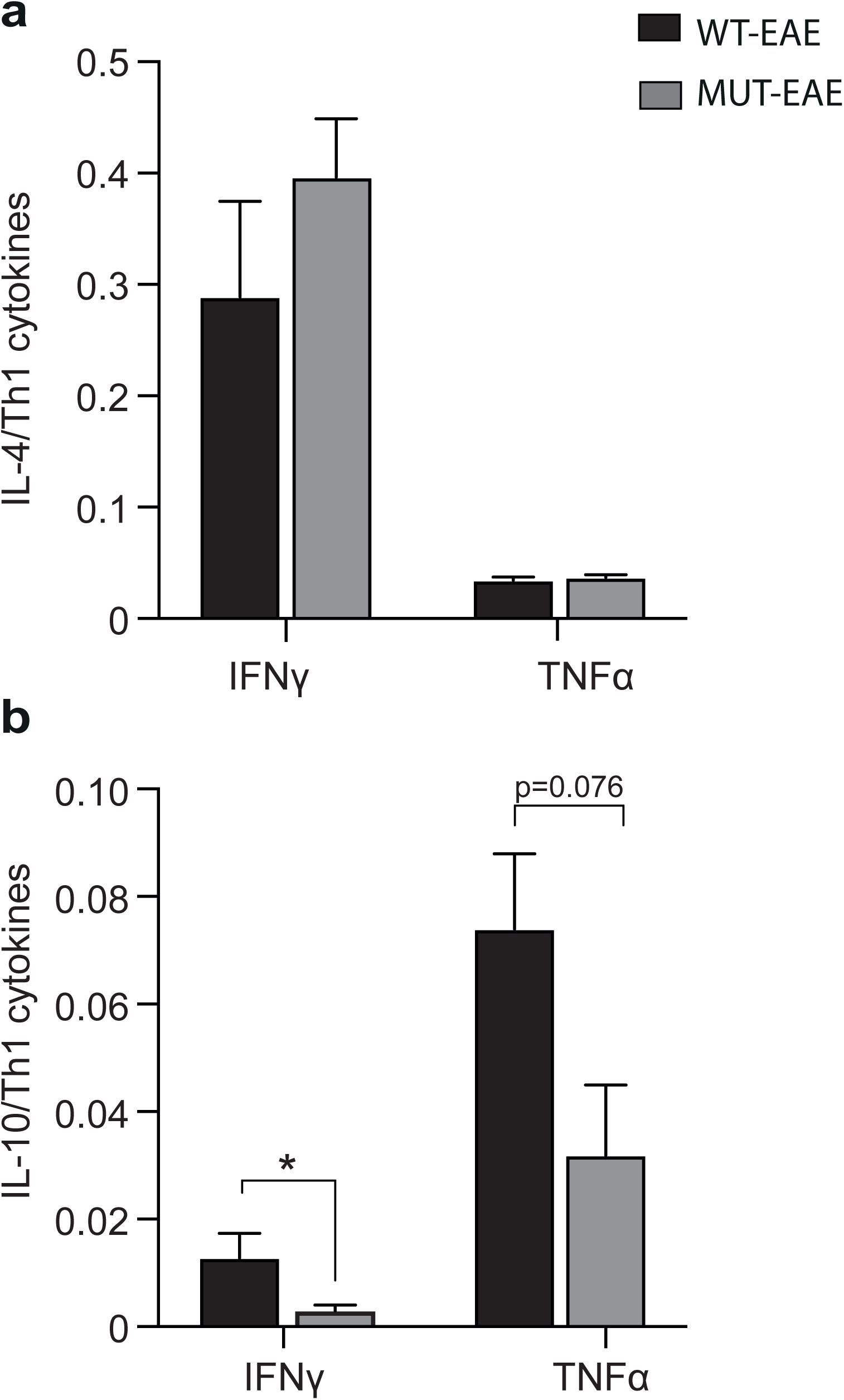
Th2/Th1 ratios during acute EAE. Comparative ratios were calculated between the production of anti-inflammatory cytokines (IL-4, IL-10) and the pro-inflammatory cytokines (IFNγ and TNFα) from WT-EAE and MUT-EAE isolated splenocytes re-stimulated *in vitro* with MOG. The ratios between **a**. IL-4/IFNγ and IL-4/TNFα, **b**. IL-10/ IFNγ and IL-10/TNFα are displayed. The lower production of IL-10 from MUT-EAE +MOG splenocytes induced a decrease in the Th2/Th1 balance during the acute stage. All values are presented as media ± SEM. A one-tail unpaired t-Student test was performed. *<0.05. n = 3 per experimental group.

#### Chronic stage

no significant levels of the tested cytokines were measured in supernatants from CFA splenocytes, irrespective of their genotype and re-stimulation with MOG (data not shown). In addition, concentration ranges for IL-2, IL-4 and IL-10 were below the detection limit of our kit, for all tested conditions.

Since IL-2 levels were undetectable in splenocytes isolated from animals in EAE chronic stage, we set to evaluate the proliferative capacity of WT and MUT splenocytes to MOG, through the incorporation of [^3^H] Thymidine, and the stimulation index (SI) was calculated per each experimental group. We found a significant SI increase in EAE splenocytes compared with CFA, regardless the genotype. In addition, no significant differences in the rate of proliferation were identified between WT and MUT cells under basal conditions (without MOG) (data not shown)

TNFα, IFNγ, IL6 and IL17 expression was induced at higher levels in MOG-stimulated groups compared with the controls (Fig. 4a-d); only MOG-induced IFNγ production was significantly higher in MUT-EAE than in WT-EAE splenocytes (Fig. 4a). Next, using cytokine levels data obtained from MOG-stimulated splenocytes from WT and MUT EAE mice, we calculated the percentage of TNFα and IFNγ downregulation from acute to chronic stage (Fig. 4e,f). Cells from MUT-EAE mice showed lower proportions of downregulation for both IFNγ and TNFα, indicating that these cells may contribute to maintain higher levels of both pro-inflammatory cytokines in MUT mice during the chronic phase of EAE. Altogether, our results indicate that, at the immune peripheral site, MeCP2 deficiency alters the autoreactivity to MOG by skewing the response towards a pro-inflammatory profile, at both acute and chronic stages of EAE.

**Fig. 4.**
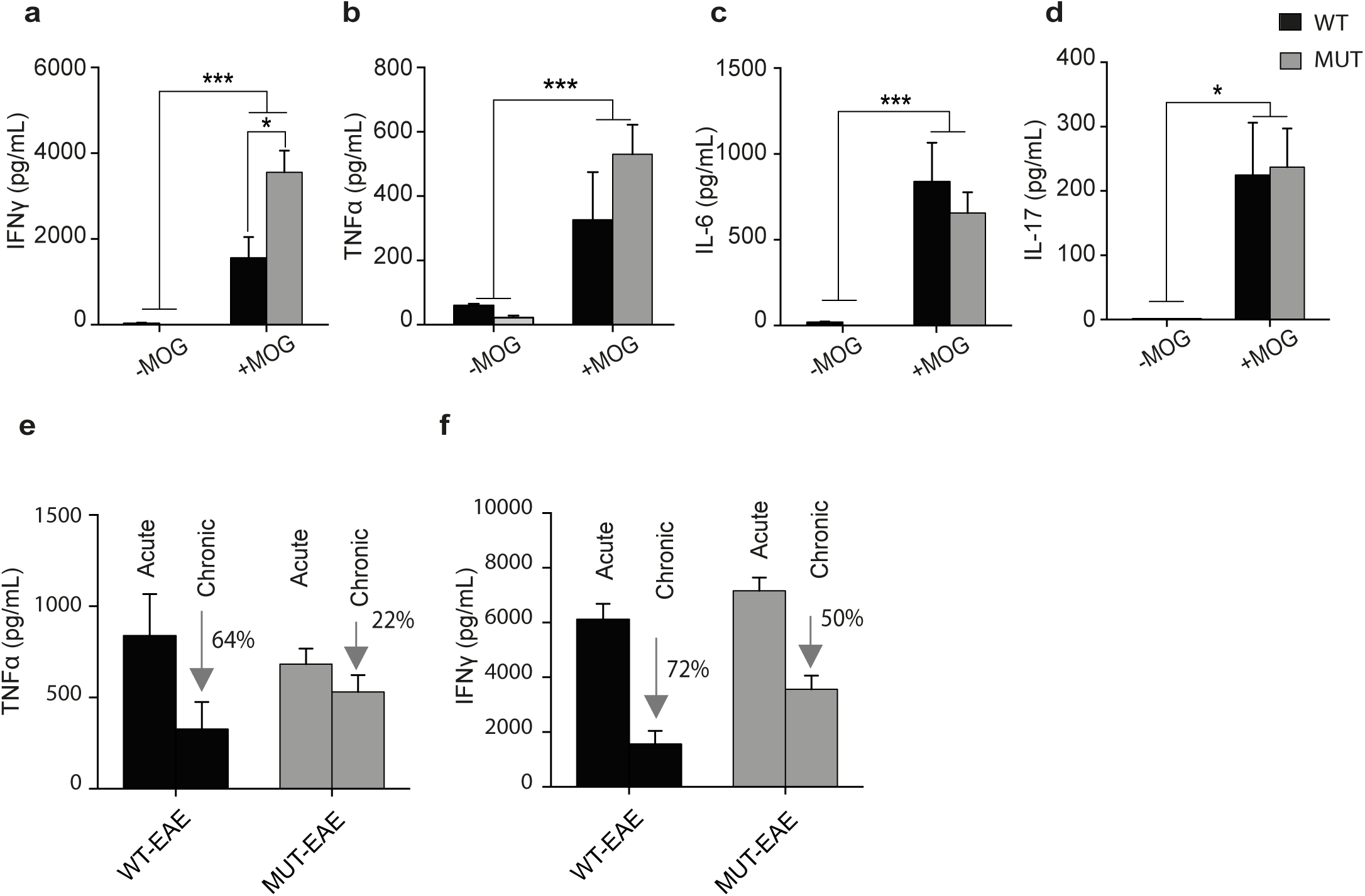
Cytokine production and proliferation of splenocytes isolated during chronic stage of EAE. This figure shows the concentration of cytokines produced by splenocytes from WT-EAE and MUT-EAE animals in the absence or presence of MOG (-MOG / +MOG) in culture for 72 hours. For all cytokines analyzed **a**. IL-6, **b**. IFNγ, **c**. TNFα and **d**. IL-17, the stimulation with MOG induced a higher and statistically significant production compared to unstimulated splenocytes. Also, IFNγ production by MUT +MOG splenocytes was significantly higher than WT +MOG. All values are presented as media ± SEM. Two-way ANOVA and Tukey post hoc test were performed when appropriate. *p<0.05; ***p<0.0002. n= 3 per experimental group. **e, f**. Regulation of TNFα and IFNγ production from acute to chronic stage of EAE. These graphs show the concentration of TNFα (e) and IFNγ (f) during the acute and chronic stages for WT and MUT splenocytes exposed to MOG. The percentage of decrease in cytokine production by WT and MUT splenocytes in the presence of MOG during the acute stage (considered as 100%) and the chronic stage was calculated. The decrease in the pro-inflammatory response of MUT splenocytes was less than that of WT when re-stimulated with MOG.

### Mecp2^308/y^ mice show higher level of infiltrating cells in spinal cord during acute EAE

Histological correlation of an autoimmune disease provides essential information on the underlying autoimmune pathology. Infiltration of T cells and mononuclear cells in the CNS is a key feature during EAE [30]. Therefore, we sought to assess whether MeCP2 deficiency could be influencing the infiltration of immune cells to the CNS, particularly at the level of the spinal cord, a target tissue in EAE. Histological analysis was performed on samples obtained during the acute and chronic stage of EAE. For this, lumbar spinal cord cryosections were stained with Toluidine Blue. To obtain the level of immune cell infiltrates in the CNS, the score corresponding to each category of infiltration was considered, from the least severe (absence of infiltrates) to the most severe (score 3, as described in methods).

No infiltrates were found in any tissue from animals treated only with CFA in both acute or chronic phase (Fig. 5a). In acute phase, lumbar spinal cord from EAE groups showed distinctive infiltration patterns (Fig.5a,b). However, MUT-EAE spinal cords (Fig.5a) showed significantly higher level of cell infiltrates in meninges and parenchyma compartments compared to WT-EAE group (Fig. 5b.i, 5b.ii). The level of infiltration of immune cells in spinal cord decreased during chronic stage, and no significant differences were found between genotypes at that time point (Fig. 5c.i, 5c.ii). These results suggest that MeCP2 mutation facilitates the infiltration of immune cells in the CNS during the acute stage of EAE.

**Fig. 5.**
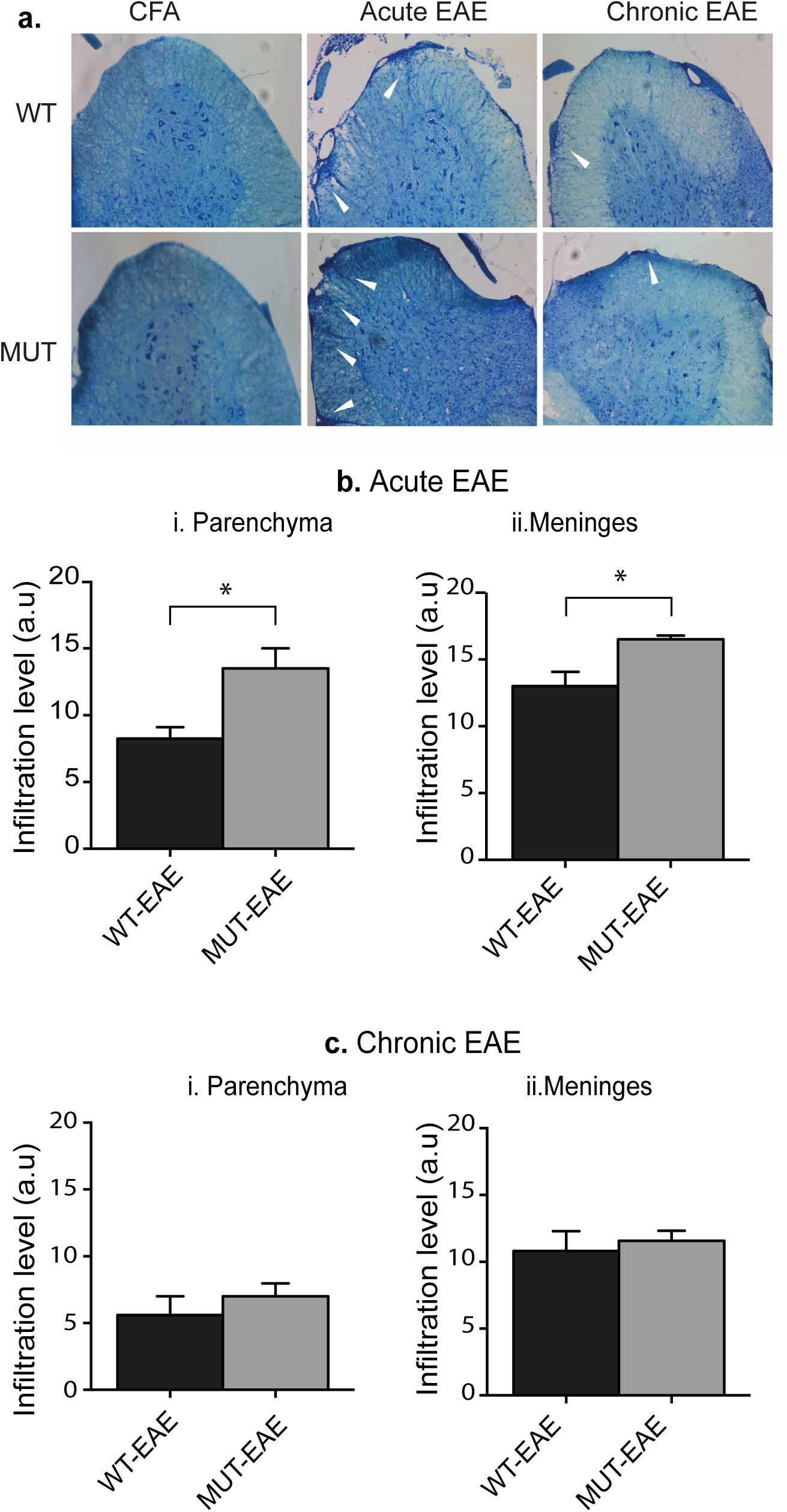
Analysis of infiltrating cells in spinal cord. **a**. Cryosections of the lumbar spinal cord were stained with Toluidine Blue and analyzed to determine the level of cellular infiltrates. Representative photos of coronal sections of spinal cords obtained from control animals (CFA, upper panel) showed no cellular infiltrates. In the lower panel, representative spinal cord images from EAE mice showed typical cellular infiltrates (white arrows). The level of infiltration was evaluated in meninges and parenchyma, considering the extent of the infiltrates and the number of cells foci. Magnification: 20X. **b-c**. During acute stage, spinal cord from MUT-EAE animals showed higher level of infiltrates in both **b**. meninges and **c**. parenchyma compared to WT-EAE. **d-e**. In chronic stage, the levels of cellular infiltrates in both spinal cord compartments were lower respect to the acute stage and with no significant differences between genotypes. All values are presented as media ± SEM. Results were analyzed using a t-Student test. *p<0.05. n= 5-7 per experimental group.

### MeCP2 deficit does not alter the expression of Iba1 and microgliosis during EAE

Myeloid cells are key for the onset and progression of EAE. These cells adopt different phenotypes that coexist in the CNS after MOG immunization, and the balance between these phenotypes and the immune mediators they release are essential for mantainance of the chronic phase [31]. Since it has been shown that MeCP2 is expressed in microglial cells and that its deletion can alter the lifespan and pathogenic phenotype in mice [32], we sought to evaluate the level of microgliosis after EAE-induction. To this end, we performed immunostainings for Iba-1 in lumbar spinal cord from all groups and quantified the Iba-1^+^ area as indicator of microgliosis during acute and chronic EAE (Fig.6a). At rest, microglial cells show a morphological phenotype characterized by long and thin processes extending from its soma. When microglia is activated, these fine processes get shorter and thicker, the cells suffer an increase in soma size, and those changes can be assessed as an increase in Iba-1^+^ area [33].

**Fig. 6.**
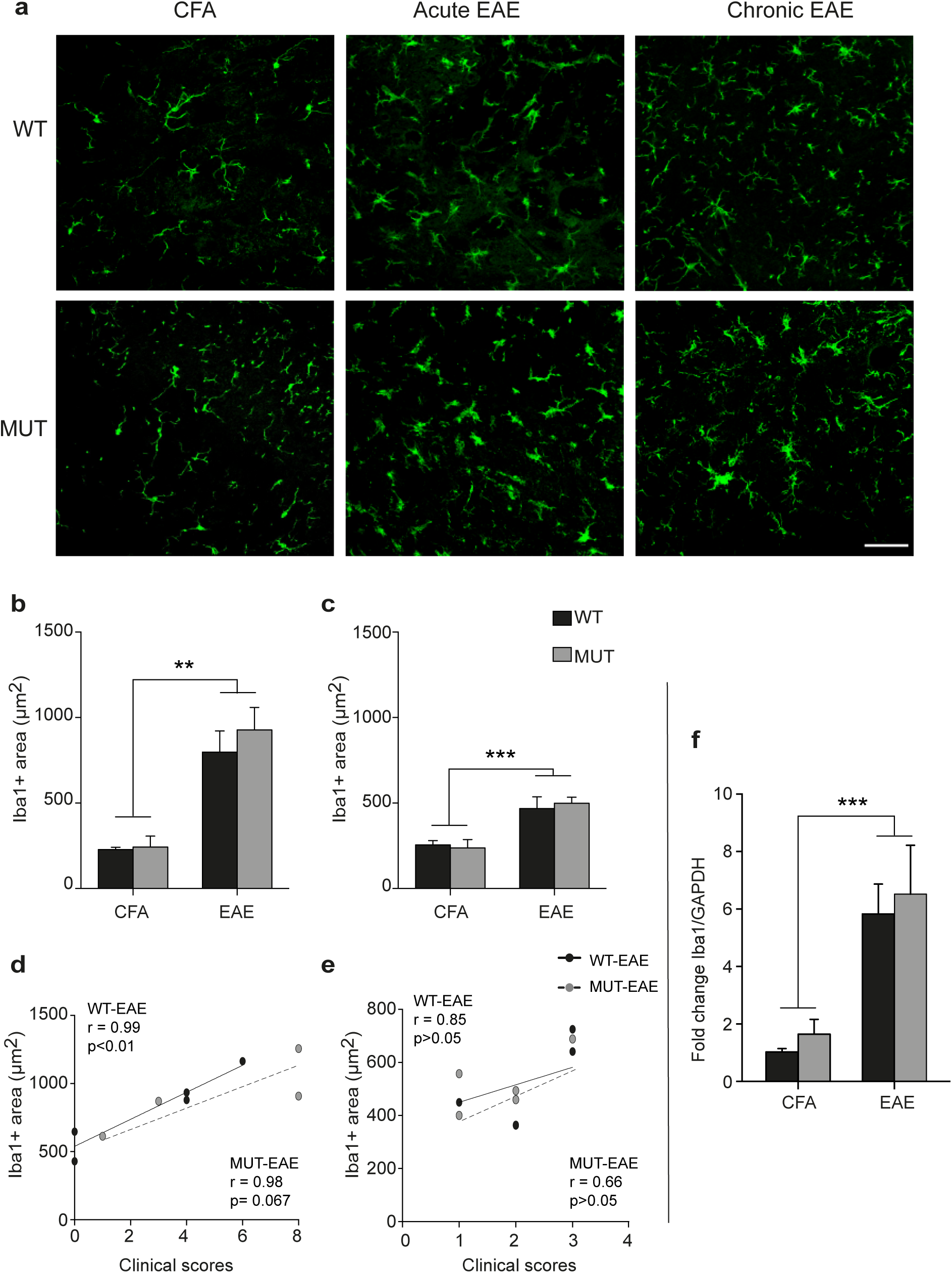
Microgliosis and Iba-1 expression analysis. **a**. Cryosections of lumbar spinal cord of all the experimental groups in acute and chronic stage can be visualized in these photographs. Microglial cells from CFA mice show small somas and thin projections. During acute EAE, cell soma increase in size, their projections get shorten and widen, which is characteristic of microgliosis and microglial activation, indicated by white arrows. In chronic stage, cells return to their “resting” morphology and others are still showing “activated” morphology. Total area occupied by Iba-1, Iba-1+ area (μm^2^), was measured by a particle size exclusion analysis. In **b**. acute and **c**. chronic stages, the Iba-1+ area was significantly higher in the EAE animal groups compared to the CFA groups. The data is presented as media ± SEM. A two-factor ANOVA was performed. **p<0.002, ***p<0.0002. **d-e**. In addition, a Pearson correlation was performed between the clinical scores and the Iba-1+ area in acute and chronic stages considering all EAE mice (WT and MUT) together. Such correlation was found to be significant between the clinical score and the Iba-1 positive area in animals in the acute stage (c) but not during the chronic stage (d). **f**. mRNA from spinal cords were isolated and through real time RT-PCR the fold change of expression was analyzed using GAPDH as a housekeeping gene. The graph shows the fold change of expression calculated as 2^-ΔΔCT^. The level of relative expression of Iba-1 during chronic stage was higher in all mice immunized with MOG compared to CFA and similar between WT and MUT mice. All values are presented as media ± SEM. Two-way ANOVA was performed. ***p<0.0021. n= 6-9 per experimental group.

Our results show that in both, acute and chronic stages, the Iba-1^+^ area (μm2) was significantly higher in EAE-induced mice (WT and MUT) compared to CFA groups (Fig. 6). No significant differences were found in the interaction between genotype and immunization with MOG in acute or chronic stage (Figure 6b,c). However, we found a significant correlation between the clinical signs evidenced at the time of euthanasia and the total area occupied by Iba1 in both WT-EAE mice and a tendency (p=0.06) in MUT-EAE groups in acute stage (Fig. 6d,e). Such correlation was not significant at chronic stage in either of EAE groups (30 dpi). To further confirm if there were subtle changes in Iba-1 gene expression, we proceeded to evaluate the levels of spinal cord mRNA isolated from all groups in chronic stage using Real Time RT-PCR. WT and MUT-EAE showed up to 5 times more levels of Iba-1 transcript than CFA groups regardless the genotypes (Fig. 6f). Similar results were obtained when we evaluated TSPO expression (translocator protein 18kDa) another marker of gliosis in EAE (12) (data not shown). Our results suggest that the deficit in MeCP2 would not affect the expression of Iba-1 and microgliosis in the context of EAE.

### Mecp2^308/y^ mice show higher and persisting inflammatory response in the CNS

The effects of infiltrating cells on the CNS during the course of EAE depend on the release of cytokines and chemokines that are responsible for neuroinflammation [31]. In order to determine the inflammatory profile at the spinal cord level during chronic stage of EAE, relative transcripts levels of several different immune mediators were determined [IFNγ, TNFα, IL-6, IL-10, IL-1β, CX3CL1 (Fractalkine), CX3CR1 (Fractalkine receptor), and FoxP3, that could provide information on the inflammatory and regulatory environment in EAE animals.

We found significant differences between CFA and EAE groups in IFNγ, TNFα, IL-10 and IL-1β mRNA expression levels (Fig. 7). In all cases, relative expression levels were significantly higher in EAE group. For IL-6 no significant difference was found between CFA and EAE groups, showing similarly low values in both cases (Fig. 7d). Importantly, MUT-EAE animals displayed higher expression of IFNγ, TNFα and IL1β compared to WT-EAE (Fig. 7a,b,c). In addition, MUT-EAE animals showed no significant increase of IL-10 mRNA respective to MUT-CFA, in opposition to WT mice (Fig. 7e).

**Fig. 7.**
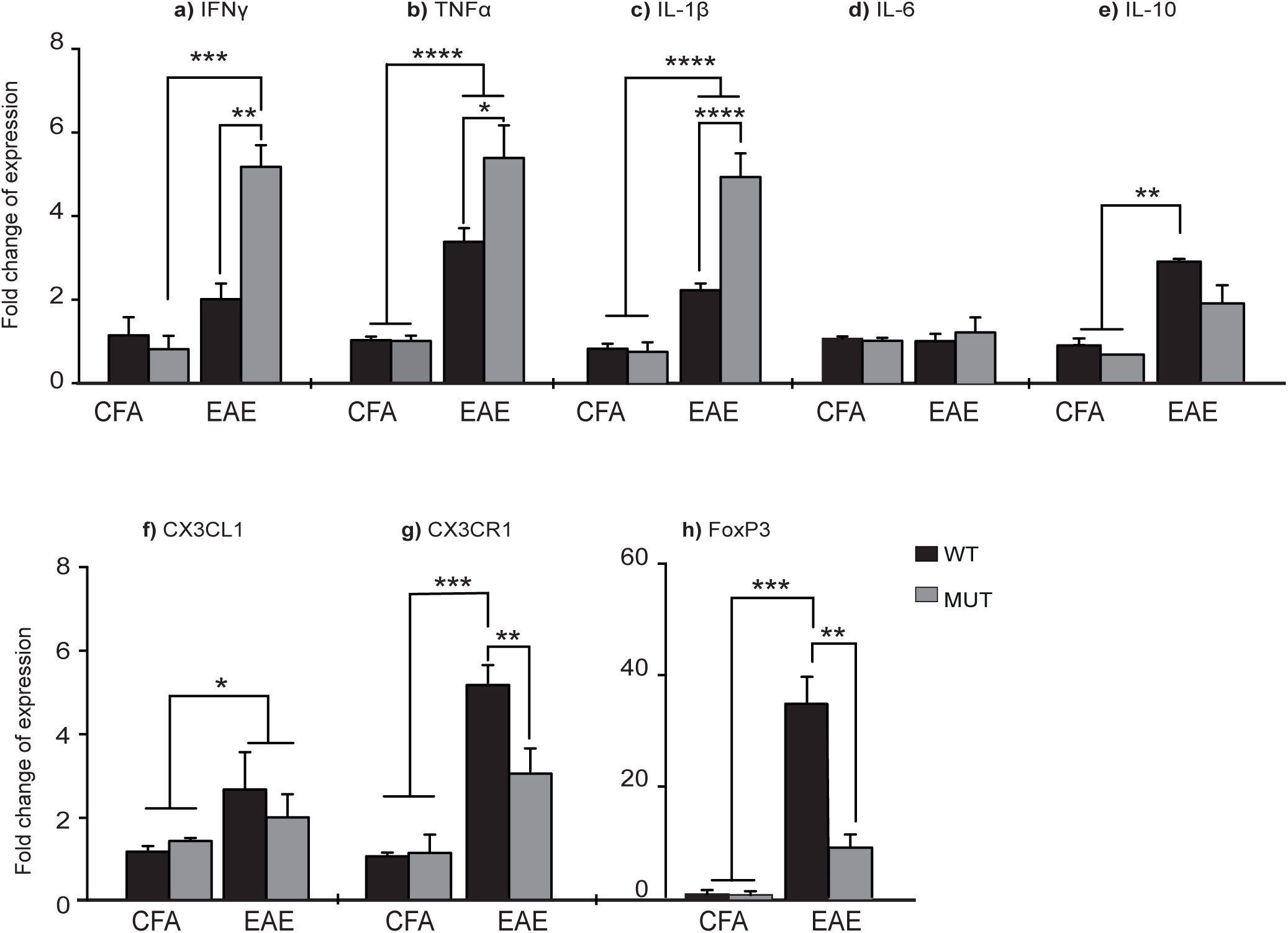
Gene expression of cytokines and immune mediators in spinal cords at chronic EAE. At 30 dpi, the lumbar spinal cords were obtained from CFA and EAE mice in order to isolate the mRNA and analyze by Real Time RT-PCR the relative level of expression of **a**. IFNγ, **b**. TNFα, **c**. IL-1β, **d**. IL-6, IL-10, **f**. CX3CL1, **g**. CX3CR1, **h**. FoxP3. Graphs show the fold change of expression calculated as 2^-ΔΔCT^. GAPDH was used as the housekeeping gene for all cases. In spinal cords from MUT-EAE, relative gene expression of IFNγ, TNFα, and IL1β (a-c) was significantly higher compared to WT-EAE. IL-10, CX3CR1 and FoxP3 gene expression (e,h,i) were significantly lower in MUT-EAE mice compared to WT-EAE, in fact, no upregulation was observed in MUT-EAE compared to CFA groups. All values are presented as media ± SEM. Two-way ANOVA was performed for each gene and Tukey post hoc test when the interaction was statistically significant. *p<0.05; **p<0.001 ***p<0.0001. n=3-6 per experimental group.

Next, we assessed the relative expression of CX3CL1 chemokine and its receptor, CX3CR1. It has been shown that the CX3CL1-CX3CR1 interaction mediates cell migration processes during EAE as well as the severity of the disease [34]. No significant differences were found in the expression of CX3CL1 between WT and MUT groups (Fig. 7f). However, the level of expression of the receptor (CX3CR1) in spinal cords from WT-EAE mice was significantly higher than in MUT-EAE (Fig. 7g). Finally, we quantified the relative expression of FoxP3, the master gene of regulatory T cells. In WT animals, EAE induction led to an increased FoxP3 expression but in MUT animals, no significant differences were found between EAE and CFA mice (Fig. 7h). Altogether these results indicate that MeCP2 deficiency promotes a persisting inflammatory profile in the CNS during an autoimmune challenge, by affecting the expression of modulatory and anti-inflammatory genes and also supporting the induction of pro-inflammatory cytokines.

## DISCUSSION

### MeCP2^308^ protein enhances autoreactivity and pro-inflammatory responses in MOG-specific cells during EAE progression

Most studies on MeCP2 and immunity have been performed in the absence of immune activation, in animal models with complete lack of MeCP2. In the present work, we set to characterize the immune response elicited by Mecp2308/y mice (MUT) subjected to EAE induction, a prototypical experimental model of neuroinflammation. Clinical signs were daily evaluated and MUT EAE mice showed higher incidence and early onset of the disease (Fig. 1), which was accompanied by an increase of infiltrating cells in spinal cord during the acute stage (Fig. 5). In this sense, an increase in CD6 has been reported in RTT patients which, in concert with its ligand ALCAM (activated leukocyte adhesion molecule), participates in leukocyte migration through the BBB [35]. Given that MUT-CFA mice did not show cellular infiltrates in the spinal cord, the increased prevalence of infiltrating cells in MUT-EAE would not be a direct consequence of alterations in the BBB permeability but rather due to an increase in autoreactive cells. In fact, MUT-EAE splenocytes, isolated during the acute stage, responded to MOG by producing higher levels of IL-2 (Fig. 2e), a cytokine that induces the clonal expansion of T cells during EAE[36], [37]. Thus, the proliferation of MOG-specific cells was boosted in a higher extent than WT-EAE mice. In addition, splenocytes from MUT-EAE mice produced higher levels of MOG-induced IL-4 in compared to WT-EAE. Although this cytokine acts as an important immunoregulatory molecule in the CNS in the context of EAE, that is not the case at peripheral immune sites [38]. Instead, lower levels of the classical immunoregulatory cytokine IL-10 were secreted by MUT-EAE cells, and IL-10/IFNγ - IL-10/TNFα ratios were reduced compared to WT-EAE group (Fig. 3b), suggesting that MUT-EAE mice generate a skewed pro-inflammatory profile in response to MOG. When we evaluated MOG-specific response in splenocytes isolated from chronic EAE mice, only IFNγ production was significantly increased in MUT-EAE compared to WT-EAE mice. However, it has been shown that the pro-inflammatory response of immune cells against myelin peptides should be downregulated to allow partial remission; this is a key feature of EAE chronic stage [31]. We showed that downregulation of TNFα expression and IFNγ were impaired in MUT-EAE splenocytes (Fig. 4 e,f). This setting resembles the immune features observed in IL-10-deficient mice [39], indicating that MeCP2 could directly regulate genes such as TNFα, IFNγ and/or IL-10, and in turn, in the absence of reciprocal regulation between these mediators, could exacerbate and maintain the pathogenesis of EAE.

Interestingly, MOG-specific response in splenocytes from either CFA and EAE animals, showed no differences in other tested cytokines between WT and MUT mice. IL-6 and IL-17 are key cytokines for EAE induction, since either IL-6- or IL-17-deficient mice are resistant to EAE [40]–[42]. Little is known about the influence of MeCP2 in the expression of IL-6 and IL-17. It has been proposed that MeCP2 is a transcriptional repressor for IL-6 gene expression [43], [44] but the exact mechanism remains unknown. MeCP2 conditional deletion in Th17 cells affects the differentiation of Th17 cells and the subsequent production of IL-17 under polarizing conditions [21]. Our present results using a mouse model of MeCP2 deficiency show a different scenario, since a decrease in Th17 cells would cause the opposite effect in EAE progression. Altogether, our results suggest that MeCP2 deficiency accelerates the onset and increase the severity of EAE by skewing the response of MOG-specific cells towards a pro-inflammatory profile, at both acute and chronic stages of EAE.

### Impaired regulation of immune mediators in the CNS of Mecp2^308/y^ mice

Since the immune profile in the CNS modulates clinical progression in EAE, we evaluated how different cytokines and immune markers are influenced by MeCP2 deficiency during chronic EAE, a time point we found greater peripheral immune deregulation (Fig. 4) and delayed motor recovery.

IL-1β is a critical mediator of EAE as mice deficient for IL-1β, NLRP3 or caspase 1 are more resistant to develop EAE[45]. Here, we showed that spinal cords from MUT-EAE mice expressed higher levels of IL-1β mRNA compared to WT-EAE, but no differences were observed at basal levels (CFA). Little is known about the role of this cytokine in disorders linked to MeCP2 mutations; however, the role of the inflammasome and its products (IL-1β and IL-18) in RTT pathogenesis has been recently explored. Pecorelli et al. demonstrated that fibroblasts isolated from RTT patients show an activated inflammasome machinery even under baseline conditions [46]. These data suggest that MeCP2 is an essential regulatory factor for the activation of the inflammasome and for preventing the perpetuation of an inflammatory state. In addition, an association between MeCP2 and IRAK1 has been described in animal models. IRAK1 is a kinase associated with the IL-1 receptor (IL-1R) that plays a central role in inflammatory responses by regulating the expression of inflammatory genes in immune cells[47]. In Mecp2-null mice it was shown that IRAK1 expression is increased in cerebellum and cortex [48]; moreover, hyper activation of IL-1 receptor pathway induces MeCP2-dependent synaptic defects linking IL-1 and immune activation to MeCP2 in neurons [49]. Therefore, it is likely that MeCP2 mutations affect IL-1β and IL-1R-mediated pathways in immune cells. Thus, this alteration may be maintained and regulated by reciprocal mechanisms involving MeCP2/IL-1β/IL-1βR/IRAK1.

Sustained production of TNFα, by infiltrating macrophages in the spinal cord is associated with increased EAE severity, promoting inflammation and preventing the establishment of the anti-inflammatory milieu needed for disease remission [50]. In the present work, MUT-EAE mice presented higher levels TNFα, expression than WT-EAE mice. High and sustained TNFα, levels are associated with pathogenic conditions that promote inflammatory processes and tissue damage [51]. In RTT patients, the analysis of the cytokine profile has shown a persistent increase in TNFα expression [19]. It is considered that MeCP2 could be a repressor for TNFα gene transcription [44], [52], [53]. However, under control conditions (CFA animals), we did not observe significant differences between the levels of TNFα expression, indicating that immune activation would be key for MeCP2 regulatory role.

Although microglial cells size and function are affected in *Mecp2*-null mice, we did not detected differences in the level of micro gliosis in WT and MUT animals (Fig. 6). Still, it is possible that specific pathways are affected by MeCP2 deficiency. CX3CL1/CX3CR1 interaction mediates communication between neurons and microglia, and it is essential to modulate basic physiological activities during development, aging and in pathological conditions [54]. Fractalkine (CX3CL1) is expressed in neurons and endothelial cells, and acts as an adhesion molecule or a soluble chemoattractant [54]. CX3CL1 signals through its CX3CR1 receptor, which is expressed primarily in microglia, monocytes/macrophages, dendritic cells, and NK [55]. Our results showed no significant differences in CX3CL1 expression between WT and MUT in spinal cord, but we found significant differences in the level of expression of CX3CR1. Increased signaling via CX3CR1 can protect against neurotoxic microgliosis; also, increased neuronal vulnerability was observed after treatment with CX3CR1 antagonists or in CX3CR1-KO mouse [54]. In Mecp2-null mouse models, CX3CR1 ablation prolonged mice lifespan, and rectified respiratory parameters and histological abnormalities in the brain [56]. However, CX3CR1 mRNA levels in hippocampus, striatum, spinal cord, cerebellum and cortex were similar in WT and Mecp2-null mice. Likewise, we found comparable expression between WT and MUT in the CFA group, but after EAE induction, CX3CR1 mRNA increased significantly only in WT mice. In this sense, EAE induction in CX3CR1-KO mice showed higher severity associated with a greater accumulation of immune cells in the CNS [34], [57]. Hence, the pathological significance of the interaction between neuron-microglia mediated by CX3CR1 may be different in pathological/physiological conditions in the context of MeCP2 mutations. In light of our results, the lower CX3CR1 expression detected in MUT-EAE mice may contribute to maintain EAE symptoms in the chronic stage.

Besides myeloid cells, MeCP2 is expressed in other immune cells, like T lymphocytes [58] and therefore, the increased EAE susceptibility observed in MUT mice could be associated with other immune populations and related cytokines. It has been reported that total loss of MeCP2 in CD4+ T cells affected cytokine-dependent activation of STAT1, a key factor for Th1 cells differentiation [21]. However, overexpression of MeCP2 in CD4+ T cells also impaired differentiation to Th1 cells, demonstrated by low levels of IFNγ production that led to an immunosuppressive phenotype[59]. Likewise, T cells from RTT patients showed decreased expression of T-bet [60], which is an essential transcription factor for IFNγ expression under activation conditions in Th1 cells [61]. This background contrasts with our results since IFNγ expression levels were significantly higher during chronic EAE in MUT animals (Fig. 7). During EAE, IFNγ acts in a stage-dependent manner: in acute stage it promotes Th1 responses and amplifies an inflammatory cascade [62]; during chronic EAE, IFNγ promotes the induction of FoxP3 and differentiation of CD4^+^ CD25^-^ T cells to regulatory T cells, that will facilitate the balance towards a Th2 profile [63], [64]. In light of our results, IFNγ in MUT mice would be exacerbating the inflammatory profile rather than promoting immunoregulation.

FoxP3 (Forkhead box P3) is a nuclear transcription factor of Treg cells that mediate tolerance and immune homeostasis [65]. The main effect of Treg FoxP3^+^ cells during EAE is to control the proliferation of autoreactive cells and the production of cytokines in the CNS [66]. During chronic stage, a higher frequency of Treg cells in the CNS correlates with EAE remission [67], [68]. FoxP3 expression is regulated by epigenetic modifications [69] and MeCP2 seems to be an important factor mediating its expression. It has been reported in Treg conditional MeCP2-KO mice, that MeCP2 is required to maintain FoxP3 expression when these cells are activated [22]. We can confirm that this effect also occurs in Mecp2^308/y^ mice, since the levels of FoxP3 expression in MUT-EAE animals were significantly lower than in WT-EAE mice (Fig. 7g). Thus, MeCP2 epigenetic regulation on Treg cells would explain in part the early onset of EAE and the presence of more severe clinical signs in MUT mice. During chronic EAE, approximately 50% of Treg cells secrete IL-10 in the CNS [68]. IL-10 is a crucial immunoregulatory cytokine and deficits in its expression lead to spontaneous autoimmunity [70]. Our results showed that MUT-EAE mice express IL-10 in the spinal cord, but at lower rates compared to WT. In fact, no significant differences were found between CFA-MUT and EAE-MUT groups (Fig. 7). Our present work resembles previous results from IL-10 KO mice, which are more susceptible to develop severe EAE and T cells exhibit a stronger specific antigen proliferation by producing higher levels of pro-inflammatory cytokines (such as TNFα and IFNγ) [39], [71]. In addition, during chronic stage, myeloid cells also change their phenotype M1 to M2 in the CNS, a crucial process associated to clinical recovery in EAE [38]. Therefore, regarding immunoregulatory processes, mutations in *Mecp2* could be affecting the release of cytokines in FoxP3+ cells, but it can also act by preventing the dedifferentiation of myeloid cells and preserving a pro-inflammatory profile in the spinal cord. Future work will evaluate the kinetics of different cell populations and immune profiles during EAE in order to understand how MeCP2 interferes with responses needed to deal with an immune insult directly in each cell population involved *in vivo*.

We can speculate that a set of immunoregulatory genes are modulated by MeCP2 in a delicate balance that will depend on the divergent characteristic of this protein. MeCP2 can either repress or stimulate gene expression depending on the context [72]. The mechanics behind these processes in the context of pro vs anti-inflammatory conditions are not fully understood. It is important to note that RTT patients do not show immunosuppression, but rather a pro-inflammatory profile [73], [74]. These discrepancies, between data reported from KO animal models and patients, are possibly related to the different degrees of loss of function of MeCP2: patients with RTT, who carry heterozygous mutations for MeCP2 in their majority, do not show large baseline immune alterations, as well as heterozygous mice [21]. Therefore, there is a wide lack of agreement along different studies concerning which genes are deregulated by Mecp2 mutations or under Mecp2-null conditions. In this sense, our work is the first study that provides an overview that begins to describe *in vivo* in real time, the complexity of neuroimmunological interactions in the context of mutant MeCP2 protein. Based on present results, we propose to design similar studies in heterozygous female mice to further identify common immune pathways affected in these models.

## Conclusions

This is the first study performed in Mecp2^308/y^ mice that explores the pathophysiology and neuroinflammation in the context of an autoimmune challenge. In light of our results we can establish that mutant MeCP2 induces an exacerbated inflammatory response in the context of EAE and provide higher susceptibility to the development of autoimmune diseases. Our results indicate that MeCP2 acts intrinsically upon immune activation, affecting neuroimmune homeostasis by regulating the pro-inflammatory/anti-inflammatory balance *in vivo*. All together, these studies may lay the groundwork to understand how the neuroimmune system maintains a pathogenic state in RTT and other MeCP2-related pathologies, and also, design effective therapeutic strategies.

## Acknowledgements

This work was supported by grants FONCyT-PICT-2013-106 and Secyt-Universidad Nacional de Córdoba, Argentina to A.L.D. MIZ, MCF and MLB were supported with fellowships financed by CONICET. ALD is a member of the Scientific Career of CONICET.

## List of Abbreviations

MeCP2: Methyl-CpG binding protein 2
RTT: Rett Syndrome
Treg: Regulatory T cells
CFA: complete Freund’s adjuvant
CNS: central nervous system
dpi: days post-induction
EAE: experimental autoimmune encephalomyelitis
GAPDH: glyceraldehyde-3-phosphate dehydrogenase
MOG: myelin oligodendrocyte glycoprotein
MS: multiple sclerosis.

